# Evaluating the efficacy of RT-qPCR SARS-CoV-2 direct approaches in comparison to RNA extraction

**DOI:** 10.1101/2020.06.10.144196

**Authors:** Ofir Israeli, Adi Beth-Din, Nir Paran, Dana Stein, Shirley Lazar, Shay Weiss, Elad Milrot, Yafit Atiya-Nasagi, Shmuel Yitzhaki, Orly Laskar, Ofir Schuster

## Abstract

SARS-CoV-2 genetic identification is based on viral RNA extraction prior to RT-qPCR assay, however recent studies support the elimination of the extraction step. Herein, we assessed the RNA extraction necessity, by comparing RT-qPCR efficacy in several direct approaches vs. the gold standard RNA extraction, in detection of SARS-CoV-2 from laboratory samples as well as clinical Oro-nasopharyngeal SARS-CoV-2 swabs. Our findings show advantage for the extraction procedure, however a direct no-buffer approach might be an alternative, since it identified up to 70% of positive clinical specimens.

## INTRODUCTION

The COVID-19 pandemic caused by SARS-CoV-2 produced significant morbidity and mortality worldwide. At the time of writing, more than six million cases and over 370,000 deaths were reported [1]. The pandemic has created an acute need for rapid, cost effective and reliable diagnostic screening. COVID-19 genetic diagnostics process include RNA extraction from Oro-nasopharyngeal swabs followed by reverse transcriptase quantitative PCR (RT-qPCR) targeting viral genes [2]. However, the global demand for reagents has placed extensive strain on supply chains for RT-qPCR kits and to an even greater extent, on RNA isolation reagents. Potentially, eliminating RNA extraction would greatly simplify the diagnostic procedure, reducing both cost and time to answer, while allowing testing to continue in case of reagent shortages. Previous studies demonstrated that several lysis buffers might allow the elimination of RNA extraction [3-5]. Very recently, two studies [6-7] used a direct no-buffer RT-qPCR approach which identified >90% of the tested clinical samples.

In this study, we tested the diagnostic efficiency following thermal inactivation (65°C for 30min and 95°C for 10min) without addition of lysis buffers (“no buffer”) or following lysis by three buffers (Virotype, QuickExtract and 2% Triton-X-100) and compared it to diagnosis after standard RNA extraction. Samples included buffers spiked with SARS-CoV-2, at concentrations 0.1-100,000 PFU/ml and 30 clinical samples, previously diagnosed as positive (20) and negative (10).

## METHODS

### RNA standards and clinical Samples

Viral RNA standards were viable SARS-CoV-2 (GISAID accession EPI_ISL_406862), cultured in Vero E6 cells and diluted in viral transport medium (Biological industries). Virus concentrations are given as plaque forming units/ml. One PFU was determined as 1000 virions by digital PCR [data not shown]. Oro-nasopharyngeal swab samples for the study were selected after approval by conventional RT-qPCR. Positive and negative samples were randomly selected for this study, kept at 4°C until use.

### RNA extraction

RNA was extracted from 200µl sample using RNAdvance Viral kit and the Biomek i7 Automated Workstation (Beckman Coulter) and eluted with 50µl H_2_O.

### Direct detection

Samples were analyzed directly or mixed 1:1 with one of the following buffers: Quick Extract DNA Extraction Solution (Lucigen), Virotype Tissue Lysis Reagent (INDICAL BIOSCIENCE GmbH) and 2% Triton-X-100 (Sigma) after inactivation at 95°C for 10min or 65°C for 30min.

### RT-qPCR

RT-qPCR assays were performed using the SensiFAST Probe Lo-ROX One-Step kit (Bioline). Primers and probe for SARS-CoV-2 E gene were taken from the Berlin protocol [2].

## RESULTS AND DISCUSSION

### SARS-CoV-2 samples in different concentrations

Standard samples were analyzed in duplicates and the results shown in figure 1 are averages. The samples were analyzed following two inactivation temperatures: 95°C for 10min or 65°C for 30min. The maximal standard deviation was <2 Ct’s with an average standard deviation of 0.4 across all samples. The limit of detection was 1 PFU/ml: In this concentration samples in the no buffer mode and Virotype at 95°C were not detected, while the RNA extraction mode averaged the lowest critical threshold (Ct=29.8) followed by QuickExtract and Triton. In 10 PFU/ml only the no buffer mode at 95°C failed to detect. The RNA extraction mode maintained the lowest Ct values across all the analyzed concentrations. The minimal delta Ct average to the RNA extraction mode was obtained using QuickExtract, followed by Triton, Virotype and the no buffer mode.

**Figure 1:**
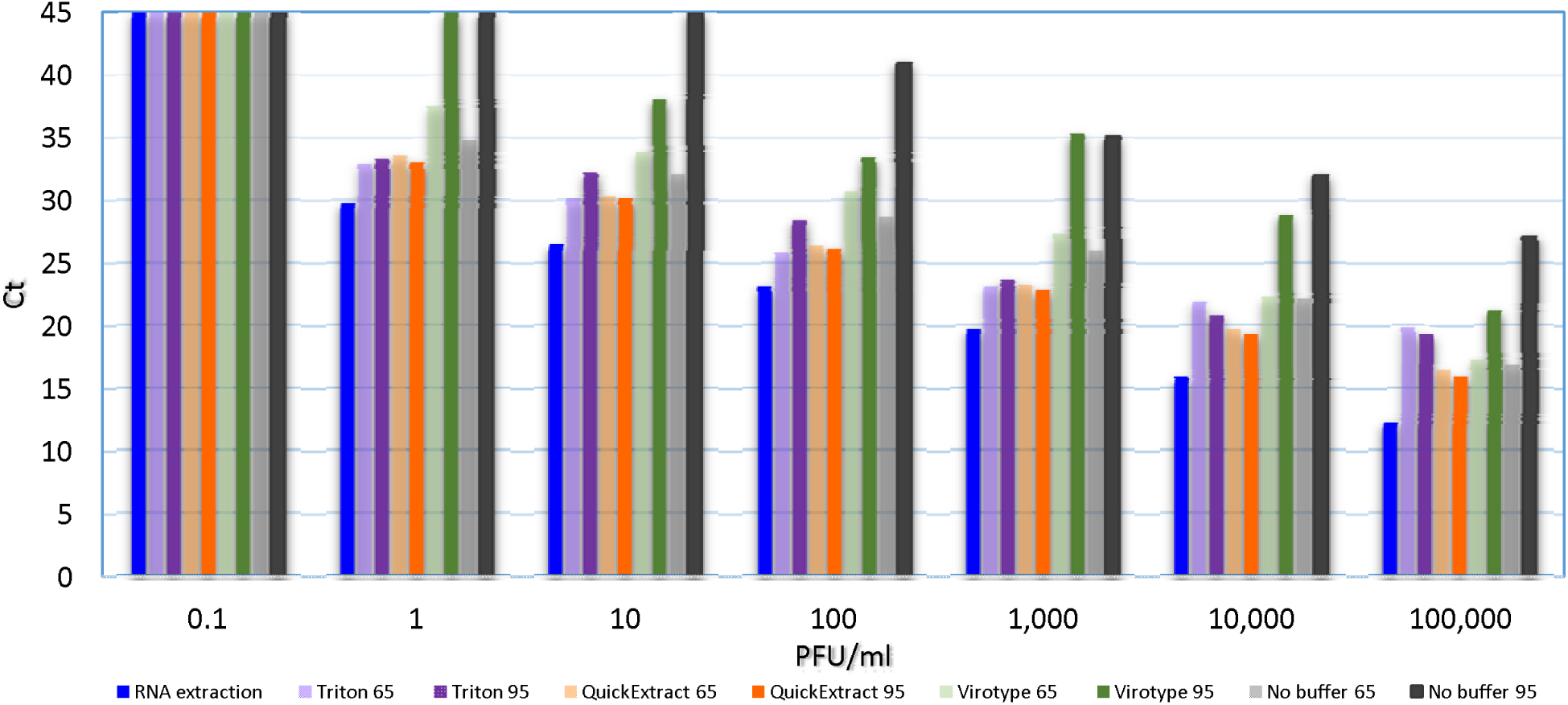
RT-qPCR results of SARS-CoV-2 samples at different concentrations. All the samples were analyzed in duplicates. Shown are averages Ct values. The different buffers and conditions are elaborated in the figure legend. Ct 45=undetected. 65=65°C, 95=95 °C.

### SARS-CoV-2 clinical samples

Next we tested the feasibility of direct SARS-CoV-2 detection in clinical samples. Twenty positive and 10 negative samples were analyzed following thermal inactivation. All previously defined negative samples remained negative across the different buffers and test conditions.

Positive samples exhibited major differences in detection capability (Table 1). The alternative buffers exhibited much lower detection levels: Triton (both inactivation protocols) detected a single positive sample (5% detection); OuickExtract and Virotype had 35-40% detection rates (both inactivation protocols). Surprisingly, direct no-buffer approach was superior with 50% and 70% for the 65°C and 95°C inactivation protocols, respectively. Detection was reversely-correlated to samples’ Ct, with efficiency dropping from 100% for Ct ≤ 32 to 25% for samples with higher Ct.

**Table 1:** RT-qPCR results of SARS-CoV-2 clinical samples using different treatments. Shown are Ct values. RNA ext. =RNA extraction. Un =undetermined (Ct = 45). Δ =the difference in Ct between RNA extraction and the different treatments. Detection level =the percentage of positive samples (Ct<45).

| Patient # | RNA ext. | Triton-X-100 2% |  |  |  | QuickExtract |  |  |  | VIROTYPE |  |  |  | No buffer |  |  |  |
| --- | --- | --- | --- | --- | --- | --- | --- | --- | --- | --- | --- | --- | --- | --- | --- | --- | --- |
|  |  | 65°C | Δ | 95°C | Δ | 65°C | Δ | 95°C | Δ | 65°C | Δ | 95°C | Δ | 65°C | Δ | 95°C | Δ |
| 1 | 17.0 | 31.9 | 15 | 31.3 | 14 | 18.6 | 1.5 | 20.4 | 3.4 | 23.7 | 6.6 | 24.2 | 7.1 | 18.3 | 1.2 | 21.9 | 4.8 |
| 2 | 19.5 | un | - | un | - | 24.4 | 4.9 | 25.4 | 5.9 | 27.9 | 8.4 | 24.9 | 5.5 | 25.4 | 5.9 | 22.7 | 3.2 |
| 3 | 21.7 | un | - | un | - | 31.4 | 9.8 | 30.9 | 9.3 | 33.8 | 12 | 30.5 | 8.9 | 28.5 | 6.8 | 27.9 | 6.3 |
| 4 | 28.8 | un | - | un | - | 33.7 | 4.9 | 35.6 | 6.9 | 35.3 | 6.5 | 33.1 | 4.3 | 31.8 | 3 | 31.7 | 3 |
| 5 | 29.0 | un | - | un | - | 33.7 | 4.7 | 35.4 | 6.4 | 38.7 | 9.8 | 35 | 6 | 32 | 3.1 | 32.5 | 3.6 |
| 6 | 29.6 | un | - | un | - | 34 | 4.4 | un | - | 35.6 | 6 | un | - | 31.4 | 1.8 | 29.7 | 0.1 |
| 7 | 30.3 | un | - | un | - | un | - | un | - | un | - | un | - | 39.1 | 8.8 | 35 | 4.7 |
| 8 | 30.3 | un | - | un | - | un | - | un | - | un | - | un | - | un | - | 38.8 | 8.6 |
| 9 | 31.2 | un | - | un | - | 36.2 | 5 | un | - | un | - | un | - | 35.2 | 4 | 35.3 | 4.1 |
| 10 | 31.4 | un | - | un | - | un | - | 42.4 | 11 | un | - | 38.4 | 7 | un | - | 36.1 | 4.8 |
| 11 | 31.6 | un | - | un | - | un | - | un | - | un | - | un | - | un | - | 35.8 | 4.2 |
| 12 | 32.0 | un | - | un | - | un | - | un | - | un | - | un | - | un | - | 37.6 | 5.6 |
| 13 | 32.4 | un | - | un | - | un | - | 37.5 | 5.1 | 35.7 | 3.3 | 38.6 | 6.2 | 35.6 | 3.2 | 36.9 | 4.5 |
| 14 | 32.9 | un | - | un | - | un | - | un | - | un | - | un | - | 38.3 | 5.4 | un |  |
| 15 | 33.4 | un | - | un | - | un | - | un | - | un | - | 38.3 | 4.9 | un | - | 34.9 | 1.5 |
| 16 | 33.7 | un | - | un | - | un | - | un | - | un | - | un | - | un | - | un | - |
| 17 | 33.8 | un | - | un | - | un | - | un | - | un | - | un | - | un | - | un | - |
| 18 | 35.7 | un | - | un | - | un | - | 39.7 | 4 | un | - | un | - | un | - | un | - |
| 19 | 35.7 | un | - | un | - | un | - | un | - | un | - | un | - | un | - | un | - |
| 20 | 35.9 | un | - | un | - | un | - | un | - | un | - | un | - | un | - | un | - |
| Detection level (%) |  | 5 |  | 5 |  | 35 |  | 40 |  | 35 |  | 40 |  | 50 |  | 70 |  |

## CONCLUSION

Our results demonstrate that RNA extraction significantly improves comprehensive and sensitive clinical diagnosis of SARS-CoV-2. We suggest that clinical samples (which include a multitude of nucleic acids and proteins) might significantly hamper detection. Although being previously reported to facilitate viral detection [3-5], the tested buffers severely compromised the limit of detection (to a maximum of 40%). This is surprising, considering that direct analysis without adding buffers achieved a 70% detection level. This no-buffer direct approach could potentially be used with some success in times of need to achieve screening for high-titer samples.

## ETHICAL STATEMENT

Pre-existing samples were used and de-identified. This work was therefore determined to be “not human subjects’ research”.

## ACKNOLEDGMENTS

SARS-CoV-2 was kindly provided by Bundeswehr Institute of Microbiology, Munich, Germany. The authors would like to thank Itai Glinert for his fruitful reviewing of this manuscript.

## REFERENCES

1. https://www.worldometers.info/coronavirus/

2. Corman VM, Landt O, Kaiser M, Molenkamp R, Meijer A, Chu DK, Bleicker T, Brünink S, Schneider J, Schmidt ML, Mulders DG, Haagmans BL, van der Veer B, van den Brink S, Wijsman L, Goderski G, Romette JL, Ellis J, Zambon M, Peiris M, Goossens H, Reusken C, Koopmans MP, Drosten C. Detection of 2019 novel coronavirus (2019-nCoV) by real-time RT-PCR. Euro Surveill. 2020 Jan;25(3):2000045. doi: 10.2807/1560-7917.ES.2020.25.3.2000045.

3. Ladha A, Joung J, Abudayyeh O, Gootenberg J, Zhang F. 2020. A 5-min RNA preparation method for COVID-19 detection with RT-qPCR. medRxiv. 2020.05.07.20055947

4. Joel D. Pearson, Daniel Trcka, Sharon J. Hyduk, Marie-Ming Aynaud, J. Javier Hernández, Filippos Peidis, Suying Lu, Kin Chan, Jim Woodgett, Tony Mazzulli, Liliana Attisano, Laurence Pelletier, Myron I. Cybulsky, Jeffrey L. Wrana, Rod Bremnera. Comparison of SARS-CoV-2 Indirect and Direct Detection Methods. doi: https://doi.org/10.1101/2020.05.12.092387. BIORXIV

5. Merindol N, Pépin G, Marchand C, Rheault M, Peterson C, Poirier A, Houle C, Germain H, Danylo A.J. SARS-CoV-2 detection by direct rRT-PCR without RNA extraction. Clin Virol. 2020 May 7;128:104423. doi: 10.1016/j.jcv.2020.104423. Online ahead of print.PMID: 32416598 Free PMC article.

6. Fomsgaard AS, Rosenstierne MW. 2020. An alternative workflow for molecular detection of 357 SARS-CoV-2 - escape from the NA extraction kit-shortage, Copenhagen, Denmark, March 358 2020. Euro Surveill 25.

7. Emily A. Bruce, Meei-Li Huang, Garrett A. Perchetti, Scott Tighe, Pheobe Laaguiby, Jessica J. Hoffman, Diana L. Gerrard, Arun K. Nalla, Yulun Wei, Alexander L. Greninger, Sean A. Diehl, David J. Shirley, Debra G. B. Leonard, Christopher D. Huston, Beth D. Kirkpatrick, Julie A. Dragon, Jessica W. Crothers, Keith R. Jerome, Jason W. Botten. Direct RT-qPCR detection of SARS-CoV-2 RNA from patient nasopharyngeal swabs without an RNA extraction step. BIORXIV. doi: https://doi.org/10.1101/2020.03.20.001008

